# Food supplementation affects gut microbiota and immunological resistance to parasites in a wild bird species

**DOI:** 10.1101/770644

**Authors:** Sarah A. Knutie

## Abstract

1. Supplemental feeding can increase the overall health of animals but also can have varying consequences for animals dealing with parasites. Furthermore, the mechanism mediating the effect of food supplementation on host-parasite interactions remains poorly understood.
2. The goal of the study was to determine the effect of food supplementation on host defenses against parasitic nest flies and whether host gut microbiota, which can affect immunity, potentially mediates these relationships. In a fully crossed design, I experimentally manipulated the abundance of parasitic nest flies (*Protocalliphora sialia*) and food availability then characterized the gut microbiota, immune responses, and nest parasite abundance of nestling eastern bluebirds (*Sialia sialis*).
3. Food supplemented birds had 75% fewer parasites than unsupplemented birds. Parasite abundance decreased throughout the breeding season for unsupplemented birds, but abundance did not change throughout the season for supplemented birds. Food supplementation increased overall fledging success. Parasitism had a sublethal effects on blood loss, but food supplementation mitigated these effects by increasing parasite resistance via the nestling IgY antibody response.
4. Food supplementation increased the gut bacterial diversity in nestlings, which was negatively related to parasite abundance. Food supplementation also increased the relative abundance of *Clostridium* spp. in nestlings, which was positively related to their antibody response and negatively related to parasite abundance.
5. *Synthesis and applications*. Overall, these results suggest that food supplementation, especially early in the breeding season, increases resistance to parasitism during the early life stage of the host, which could be mediated by the effect of supplementation on the gut microbiota. Wildlife food supplementation is a common pastime for humans worldwide and therefore it is important to determine the consequences of this activity on animal health. Furthermore, supplemental feeding could induce resistance to detrimental parasites (e.g. invasive parasites) in hosts when management of the parasite is not immediately possible.

## 1 INTRODUCTION

Environmental factors, such a food availability, can influence host-parasite interactions (Becker, Streicker & Altizer 2015; Becker *et al*. 2018; Sánchez *et al*. 2018). Host defense strategies against parasites, such as tolerance and resistance, are often condition-dependent and regulated by resource availability (Lee *et al*. 2006; Cotter *et al*. 2011; Sternberg *et al*. 2012; Howick & Lazzaro 2014; Knutie *et al*. 2017c). Tolerance mechanisms, such as tissue repair or compensation for energy loss, reduce the damage that parasites cause without reducing parasite fitness (Miller, White & Boots 2006; Råberg, Sim & Read 2007; Read, Graham & Råberg 2008; Medzhitov, Schneider & Soares 2012). For example, avian parents from parasite-infested nests reduce the cost of parasitism by feeding their offspring more than parents from non-parasitized nests (Christe et al. 1996; Tripet and Richner 1997; Knutie et al. 2016). Consequently, despite increasing parasite loads, offspring do not suffer a high cost of parasitism because they are able to compensate for resources lost to the parasites.

Another condition-dependent defense mechanism is resistance, such as the immune response, which reduces the damage that parasites cause by reducing parasite fitness (Read *et al*. 2008). Resistance can be condition-dependent because mounting immune responses can be energetically costly and therefore only hosts with enough food resources may be able to invest in immunity (Sheldon & Verhulst 1996; Svensson *et al*. 1998; Lochmiller & Deerenberg 2000; Demas 2004; Sternberg *et al*. 2012; Howick & Lazzaro 2014; Cornet *et al*. 2014). One explanation for the positive relationship between food availability and immunity is that the extra nutrients directly increase the production of immune cells (Strandin, Babayan & Forbes 2018). For example, supplemented protein can increase the concentration of cellular immune cells (e.g. eosinophils, globule leukocytes and mast cells) (reviewed in Coop and Kyriazakis 2001) and humoral immune cells (e.g. Ig antibodies) (Datta *et al*. 1998).

Food availability might also indirectly influence resistance by affecting the gut microbiota of the host. Host diet composition can directly alter the gut microbiota of the host (David *et al*. 2014; Carmody *et al*. 2015). For example, hosts that consume a plant-based diet have significantly different bacterial communities than hosts that consume an animal-based diet (David *et al*. 2014; Knutie *et al*. 2017a). Since the gut microbiota can affect the development and maintenance of the immune system of the host (reviewed in Round and Mazmanian 2009; Hooper et al. 2012), diet-based shifts in the host gut microbiota might also affect parasite resistance. However, studies are still needed to determine whether the host microbiota plays a role in mediating the effect of food availability on parasite resistance.

Humans can change resource availability for animals by increasing the abundance of unnatural food using wild bird feeders and the disposal of human trash (Murray *et al*. 2016; Bosse *et al*. 2017; Start *et al*. 2018). In fact, humans provide many wild bird species with a large proportion of their food (Jones & James Reynolds 2008; Jones 2011; Cox & Gaston 2018), which can play a key role in maximizing the birds’ fitness (Tollington *et al*. 2019). In the United States alone, approximately 50 million households provide over a half a million tons of supplemental food to attract wild birds to their property (Robb *et al*. 2008; Cox & Gaston 2018). Supplemental feeding of birds can have several benefits to birds and humans. Feeding wild birds can improve the mental health of humans along with their connection with nature (Jones 2011; Cox & Gaston 2016, 2018; Cox *et al*. 2016; Shaw, Miller & Wescott 2017). Birds that are supplemented with food are often in better condition, which in turn can increase their reproductive success (Tollington *et al*. 2019) and enhance some measures of immunity (Lochmiller, Vestey & Boren 1993; Wilcoxen *et al*. 2015; Ruhs, Vézina & Karasov 2018; Sánchez *et al*. 2018; Strandin *et al*. 2018). Because supplemental feeding can increase bird abundance (Fuller *et al*. 2008), it can also increase pathogen transmission by increasing host aggregation and contact at the feeders (Wilcoxen *et al*. 2015; Becker *et al*. 2015). Although correlational studies have variably linked food availability and host-parasite outcomes, causal tests are still needed to determine the effect of food supplementation on host-parasite interactions and how the host microbiota could be affecting parasite resistance (Becker *et al*. 2015; Altizer *et al*. 2018).

The eastern bluebird (*Sialia sialis*) is a North American bird species that is unnaturally fed by humans. In the 1970s, populations of eastern bluebirds declined, which was thought to be linked to a loss of suitable foraging and nesting habitat (Gowaty & Plissner 2015). In response, humans built and established artificial nest boxes and started supplementing the birds’ natural diet of insects, spiders, and small fruits (Pinkowski 1977a) with mealworms. Since the 1970s, the eastern bluebird population size has rebounded (Sauer & Droege 1990) but humans continue to maintain nest boxes and provide bluebirds with supplemental food, such as mealworms. Bluebirds are also infested with parasitic nest flies (*Protocalliphora* sp.) throughout their range. Past studies have found highly variable blowfly abundances along with both a negative and no effect of blowflies on fledging success of bluebirds (Johnson 1929; Pinkowski 1977b; Roby, Brink & Whitmann 1992; Wittmann & Beason 1992; Smar 1996; Hannam 2006). One explanation for the variable parasite load and relationship between bluebirds and their parasites is that the bluebird populations have access to different resources (natural and supplemental), which in turn, could affect their ability to defend themselves.

The goal of this study was to determine the effect of food supplementation on host defenses against parasites, whether these effects differ across the breeding season, and whether the gut microbiota of the host could be mediating the effect of food on host defense strategies. Across the breeding season, I experimentally manipulated parasite abundance and food availability (by supplementing birds with mealworms (*Tenebrio molitor*) or not) then quantified growth, hemoglobin levels (proxy of blood loss), glucose levels (proxy of energy), and survival (fledging success) of nestling bluebirds. I also quantified the *Protocalliphora*-binding antibody response, as a measure of resistance, and the gut microbiota of nestlings.

Bluebird nestlings are naturally tolerant of *Protocalliphora sialia* in relation to growth and survival but suffered sublethal effects (blood loss) from the parasite (Grab *et al*. 2019). If food supplementation supports host tolerance without affecting resistance, then parasitized birds that are supplemented will maintain good health (i.e. growth, survival) without changes in their parasite load. However, because food supplementation can enhance the immune response of birds (Strandin *et al*. 2018), birds with supplemented food are predicted to shift their host defense strategy from tolerance to resistance in order to reduce the sublethal effect of the parasite on blood loss. Therefore, I predicted that food supplementation would reduce parasite load in the naturally parasitized control nests. Because insect (food) availability can increase throughout breeding season (Bowlin & Winkler 2004), I predicted that supplementation would be less effective at reducing parasite abundance later in the breeding season.

Food supplementation could either directly or indirectly increase host resistance to parasitism. More nutrients could provide additional resources to directly speed up immune system development, such as the IgY antibody response or haptoglobin production, in nestlings. When a host is bitten by an ectoparasite, a series of immune pathways is activated by the host to induce an inflammatory response that acts to prevent or reduce feeding by the ectoparasite (reviewed in Owen, Nelson & Clayton 2010). Briefly, tissue damage and the introduction of antigens from the parasite stimulate the release of pro-inflammatory cytokines, which can produce acute phase proteins (e.g. haptoglobins). These chemical substances signal cells of the innate immune system, such as macrophages, to travel to the damaged tissue to degrade antigens and cells of the adaptive immune response induce the production of antigen-specific antibodies (e.g. IgY antibodies), which form lasting memory cells that can be activated quickly when the host is re-exposed to the parasite (antigen). The immune cascade described above can negatively affect ectoparasites by causing edema (tissue swelling), which prevents the parasites from feeding from capillaries, and can damage the parasite’s tissue with proteolytic molecules. Consequently, these actions can reduce the fitness of the parasite by reducing blood meal size (Owen *et al*. 2009). However, an immune response can be energetically costly to produce and therefore only well-fed nestlings might be able to produce this response (Sheldon & Verhulst 1996; Lochmiller & Deerenberg 2000; Cornet *et al*. 2014).

Alternatively, diet can affect the gut microbiota (e.g. particular taxa) of the host (Knutie *et al*. 2017a), which could impact parasite resistance. Therefore, I predicted that food supplementation would increase the gut bacterial diversity of the nestlings, which would be positively correlated with the immune responses and negatively correlated with parasite abundance. Additionally, a particular species of symbiont (e.g. gut bacterial pathogens) could affect another species due to its earlier arrival by modulating the immune response (Sousa 1992; de Roode *et al*. 2005; Devevey *et al*. 2015). Therefore, I also explored the relationship between ectoparasite abundance and the relative abundance of bacterial genera that included avian pathogens, such as *Campylobacter spp., Clostridium spp., Enterococcus spp., Escherichia spp., Lactobacillus spp., Listeria spp., Mycobacteria spp., Pasteurella spp., Salmonella spp., Staphylococcus spp., Streptococcus spp*., and *Vibrios spp.* (reviewed in Benskin *et al*. 2009).

## 2 MATERIAL AND METHODS

### 2.1 Study system

From May to July 2017, approximately 150 nest boxes were monitored in Clearwater and Hubbard County in Northern Minnesota, USA, near the University of Minnesota Itasca Biological Station and Itasca State Park (47°13’33” N, - 95°11’42” W). Eastern bluebirds are abundant at the site and nest in artificial cavities. *P. sialia* is the only ectoparasite that infests the nests of bluebirds at this site.

### 2.2 Experimental manipulation of parasites and food availability

Parasite abundance and food availability were experimentally manipulated for each nest using a 2×2 factorial design. Boxes were checked once a week for nesting activity. Once eggs appeared, nests were checked every other day until nestlings hatched. When eggs were incubated for 10 days, mealworm feeders were placed 10 m from the nest box (Fig. S1), with the intention that the feeder would only provide food to the birds from this focal nest. The feeders were constructed from pine wood (2.54 cm by 10.16 cm) and the basin for the mealworms was an empty, sterilized aluminum cat food can. The can had pin sized holes in the bottom of the can to allow for drainage from the rain and the can was attached to the wood platform with Velcro. The feeders were attached to a green garden fence post with 18-gauge aluminum wire, approximately 1.5m above the ground.

Nests were assigned to the mealworm-supplemented or non-supplemented treatment. For the supplemented treatment, 15 mealworms (Rainbow Mealworms, Compton, CA) per nestling per day were added to the feeders until the bird eggs hatched, which was based on the recommended feeding regime by the North American Bluebird Society. For the non-supplemented treatment, the feeders were visited daily but mealworms were not added to the feeders to control for the effect of disturbance on the birds. Parents were supplied with mealworms a few days before nestlings hatched to increase the chances that parents would find the feeders before their nestlings hatched. Birds were supplemented or not until nestlings reached 10 days old.

Nests were also assigned to the parasitized or non-parasitized treatment when eggs hatched. The nestlings and top liner of the nest were removed in order to treat the nest with either water to allow for natural parasitism or a 1% permethrin solution (Permacap) to remove all parasites (Knutie *et al*. 2016; DeSimone *et al*. 2018). Nestlings never contacted the parasite treatment since the liquid was sprayed under the nest liner.

### 2.3 Nestling growth and survival

Within two days of hatching, nestlings (g) were weighed using a portable digital scale balance and tarsus length (mm), bill length (mm), and first primary feather length (mm) were measured using dial calipers. When nestlings were ten days old, they were measured again and banded with a numbered metal band. When nestlings were approximately 13 days old, the boxes were checked every other day from a distance (to avoid pre-mature fledging) to determine the fledging success and age at which the nestlings fledged or died (>10-day old nestlings are not typically removed from the nest by the parents after they die, S.A.K. personal obs.).

### 2.4 Sample collection

Feces and a small blood sample (<30 µL) were collected from nestlings when they were 10 days old. Hemoglobin was measured from whole blood using a HemoCue® HB +201 portable analyzer and glucose was measured using a HemoCue® Glucose 201 portable analyzer. Blood samples were placed on ice until they were centrifuged for 3 minutes at 10000 rpm to separate the plasma from the red blood cells. Plasma and red blood cells were then stored separately in a −80°C freezer. Fecal samples were placed on ice until stored in a −80°C freezer. Samples were then transported on dry ice to the University of Connecticut. Enzyme-linked immunosorbent assays (ELISA) and Tridelta PHASE haptoglobin assay (TP-801) were then performed to quantify *P. sialia*-binding antibody (IgY) and haptoglobin (acute phase protein) levels, respectively, in the plasma of parasitized nestling. IgY antibody levels from unsupplemented nestlings were first published in (Grab *et al*. 2019); see this paper for the detailed protocol for the IgY ELISA.

### 2.5 Quantifying parasites

Once nestlings died or fledged, nests were collected and stored in plastic bags. Within eight hours of collection, nests were dissected and all larvae, pupae, and pupal cases were counted to determine total parasite abundance for each nest. Eclosed flies were collected and identified as *Protocalliphora sialia*.

### 2.6 Bacterial DNA extraction and sequencing

Total DNA was extracted from nestling bluebird feces from parasitized nests and three mealworms using a MoBio PowerFecal DNA Isolation Kit. DNA extractions were then sent to the University of Connecticut Microbial Analysis, Resources and Services for sequencing with an Illumina MiSeq platform and v2 2×250 base pair kit (Illumina, Inc). A laboratory blank was also sequenced to control for kit contamination and found no detectable sequences. Bacterial inventories were conducted by amplifying the V4 region of the 16S rRNA gene using primers 515F and 806R and with Illumina adapters and dual indices (Kozich *et al*. 2013). Raw sequences were demultiplexed with onboard bcl2fastq and then processed in Mothur v1.39.5 (Schloss *et al*. 2009) according to the standard MiSeq protocol (Kozich *et al*. 2013). Briefly, forward and reverse sequences were merged. All sequences with any ambiguities, that did not align to the correct region, or that did not meet length expectations, were removed. Sequences were aligned to the Silva nr_v119 alignment (Quast *et al*. 2013). Chimeric reads were also removed using UCHIME (Edgar *et al*. 2011). Non-bacterial sequences that classified as chloroplasts, mitochondria, or unknown (i.e. did not classify to the level of kingdom) were removed. Sequences were grouped into operational taxonomic units (OTUs) based on a 97% similarity level and identification of the OTUs was done using the Ribosomal Database Project Bayesian classifier (Wang *et al*. 2007) against the Silva nr_v119 taxonomy database. Alpha (sobs, Shannon index, Simpson index) and beta diversity statistics were calculated by averaging 1,000 random subsampling of 10,000 reads per sample. The resulting data sets included a total of 702,648 sequences and an average of 41,332 reads per sample (min: 17,582, max: 57,283).

### 2.7 Statistical analyses

A generalized linear model (GLM) with zero-inflated Poisson errors (to control for the number of zeros in the fumigated treatment) was used to determine whether parasite treatment, food treatment, and their interaction affected parasite abundance. Within the sham-fumigated treatment, a negative binomial GLM was used to determine whether food supplementation, timing of breeding (Julian date), and their interaction affected parasite abundance.

A GLM with binomial errors for proportional data (i.e. logistic regression) was used to determine the effect of the food treatment, parasite treatment, and their interaction on fledging success. A GLM was also used to determine the effect of the interaction between the food treatment and parasite abundance on fledging success to test whether food treatment affected host tolerance of the parasite (Simms 2000); i.e. if the effect of the interaction on fledging success is significant then nestlings from each food treatment differ in parasite tolerance. Julian date was originally included as a covariate for the growth and fledging success models but it was excluded from all models because it did not account for a significant amount of variation.

Nestling growth metrics (bill length, tarsus length, first primary feather length, body mass) were not highly correlated (pairwise correlation coefficients ranged from 0.37 to 0.66; Table S1); therefore, individual generalized linear mixed models (GLMM) with nest as a random effect were used to determine the effect of parasite treatment, food treatment, and their interaction on each log_10_ transformed nestling growth metrics. GLMMs with nest as a random effect were also used to determine the effect of parasite treatment, food treatment, and their interaction on log_10_ transformed hemoglobin levels (proxy of blood loss) and glucose levels (proxy of energy levels).

Within the sham-fumigated treatment, GLMMs with nest as a random effect were used to determine the effect of food treatment on immune responses (antibody levels) and alpha bacterial diversity metrics (Shannon index and the log_10_ of sobs and Simpson index). A GLM was used to analyze the effect of food treatment on haptoglobin levels because haptoglobin was measured in only one nestling per nest. Analyses were conducted using the glm (GLM) and glmer (GLMM) functions with the lme4 package or glmmTMB (zero-inflated Poisson GLM) function with the glmmTMB package. Probability values were calculated using log-likelihood ratio tests using the Anova function in the car package (Fox & Weisberg 2002).

Microbiota data were also analyzed from mealworm samples and parasitized nestlings from both food treatments. The effect of food treatment on bacterial community dynamics in parasitized nestlings was analyzed with the Bray-Curtis Dissimilarity Matrices using PERMANOVA+ (2008, version 1.0.1; with 999 permutations) in PRIMER (2008, version 6.1.11). Relative abundances (arcsine square root transformed; Shchipkova et al. 2010; Kumar et al. 2012) of bacterial phyla and genera of birds and mealworms were analyzed using ANOVAs with food treatment and mealworms as independent variables; false discovery rate (FDR) tests were used to control for multiple analyses and Tukey post-hoc tests were used for pairwise interactions. Relative abundances of six bacterial genera that include pathogenic species were also compared between food treatments ANOVAs with FDR corrections. Analyses were conducted in R (2017, version 3.4.3). All figures were created in Prism (2017, version 7).

## 3 RESULTS

### 3.1 Effect of treatment and timing of breeding on parasite abundance

Fumigation of nests with permethrin was effective at reducing parasite abundance to zero (Fig. 1A) (Table S2) (GLM, χ^2^ = 29.83, *df* = 1, *P* < 0.0001). Food supplementation and the interaction between food and parasite treatments also affected parasite abundance (Fig. 1A) (GLM, food: χ^2^ = 12.88, *df* = 1, *P* < 0.001; interaction: χ^2^ = 9.37, *df* = 1, *P* = 0.002). Food supplementation decreased parasite abundance within the sham-fumigated treatment: 100% (11/11) of unsupplemented nests and 55.6% (5/9) of supplemented were infested with parasites and unsupplemented nests had, on average, four times as many parasites as supplemented nests (Fig. 1A). Overall, parasite abundance decreased with timing of breeding (Table S3) (Julian date: χ^2^ = 0.20, *df* = 1, *P* = 0.66) but more specifically, there was an interaction between food treatment and Julian date (interaction: χ^2^ =4.18, *df* = 1, *P* = 0.04); parasite abundance decreased throughout the season in unsupplemented nests, but not supplemented nests (Fig. 1B).

**Fig. 1.**
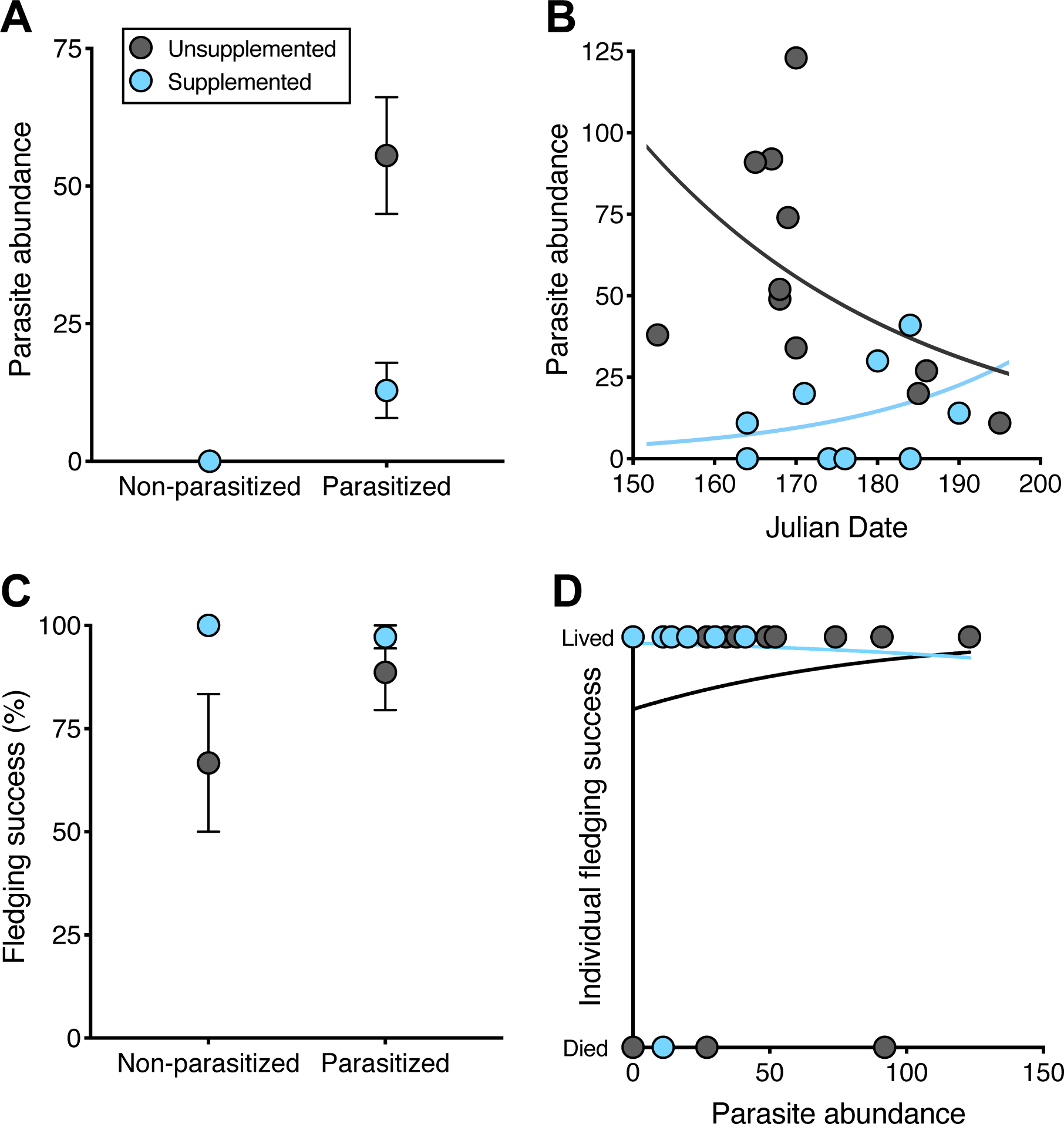
Effect of food and parasite treatment on parasite load and fledging success of bluebirds. Birds that were supplemented with food were more resistant to parasites than birds that were not supplemented (A). Parasite abundance decreased throughout the breeding season in the unsupplemented treatment but not the supplemented treatment (B). Parasitism did not affect fledging success, but supplemented birds had marginally higher fledging success than unsupplemented bird (C). Within the parasitized treatment, birds from each treatment were tolerant to their respective parasite abundances (D).

### 3.2 Fledging success and nestling growth

Parasite and food treatment affected fledging success (GLM, parasite: χ^2^ = 4.09, *df* = 1, *P* = 0.04; food: χ^2^ = 6.78, *df* = 1, *P* = 0.009), with sham-fumigated nests having higher fledging success than fumigated nests and supplemented nests having higher fledging success than unsupplemented nests (Fig. 1C). However, the interaction between parasite and food treatment did not affect fledging success (χ^2^ = 2.35, *df* = 1, *P* = 0.13). The relationship between fledging success and parasite abundance did not differ across food treatments (Fig. 1D) (i.e. tolerance to parasitism did not differ; χ^2^ = 0.10, *df* = 1, *P* = 0.75).

Parasite treatment, food treatment, and the interaction between the two factors did not significantly affect bill length (Tables 1 and S4) (GLMM, parasite: χ^2^ = 0.69, *df* = 1, *P* = 0.41, food: χ^2^ = 0.99, *df* = 1, *P* = 0.32, interaction: χ^2^ = 0.23, *df* = 1, *P* = 0.63). Parasite treatment had a marginally non-significant effect on tarsus length (GLMM, parasite: χ^2^ = 3.65, *df* = 1, *P* = 0.06), but food treatment and the interaction between the treatments did not affect tarsus length (food: χ^2^ = 0.99, *df* = 1, *P* = 0.32, interaction: χ^2^ = 0.72, *df* = 1, *P* = 0.40). Food treatment affected nestling body mass and first primary feather length (mass: χ^2^ = 5.09, *df* = 1, *P* = 0.02; feather: χ^2^ = 3.36, *df* = 1, *P* = 0.07) with unsupplemented nestlings having lower body mass and slower feather growth than supplemented nestlings (Table 1). Parasite treatment and the interaction between parasite and food treatment did not affect body mass and first primary feather length (Table 1) (mass: parasite, χ^2^ = 0.04, *df* = 1, *P* = 0.83; interaction, χ^2^ = 0.01, *df* = 1, *P* = 0.92; feather: parasite, χ^2^ = 1.13, *df* = 1, *P* = 0.29; interaction, χ^2^ = 0.38, *df* = 1, *P* = 0.54).

**Table 1.** Effect of parasite and food treatment on nestling growth, physiology, and fledging success. Numbers are mean ± SE and numbers in parentheses are the number of nests.

| Measurement | Non-supplemented |  | Supplemented |  |
| --- | --- | --- | --- | --- |
|  | Non-parasitized | Parasitized | Non-parasitized | Parasitized |
| Parasite | 0.00 ± 0.00 | 55.55 ± 10.60 | 0.00 ± 0.00 | 12.89 ± 5.00 |
| abundance | (9) | (11) | (6) | (9) |
| Bill length (mm) | 5.00 ± 0.12 | 5.19 ± 0.14 | 5.26 ± 0.22 | 5.27 ± 0.10 |
|  | (6) | (11) | (6) | (9) |
| Tarsus length | 18.35 ± 0.19 | 18.60 ± 0.23 | 18.32 ± 0.25 | 18.93 ± 0.13 |
| (mm) | (6) | (11) | (6) | (9) |
| 1<sup>st</sup> primary length | 13.64 ± 1.06 | 16.41 ± 1.51 | 17.26 ± 1.43 | 17.81 ± 0.81 |
| (mm) | (6) | (11) | (6) | (9) |
| Mass (g) | 25.71 ± 0.77 | 25.50 ± 0.77 | 27.21 ± 0.52 | 27.11 ± 0.56 |
|  | (6) | (11) | (6) | (9) |
| Hemoglobin | 11.21 ± 0.47 | 8.29 ± 0.83 | 11.55 ± 0.30 | 10.54 ± 0.88 |
| levels (g/dL) | (5) | (11) | (6) | (9) |
| Glucose levels | 282.50 ± 17.53 | 287.30 ± 32.50 | 321.30 ± 25.09 | 293.40 ± 18.80 |
| (mg/dL) | (5) | (11) | (6) | (9) |
| Nestlings fledged | 26/34 (77%) | 48/51 (94%) | 21/21 (100%) | 38/39 (97%) |

### 3.3 Nestling hemoglobin and glucose levels

Parasite treatment affected hemoglobin levels (Table S4) (GLMM, χ^2^ = 5.55, *df* = 1, *P* = 0.02) with parasitized nestlings having lower hemoglobin than non-parasitized nestlings (Table 1). Food treatment had a marginally non-significant effect on hemoglobin levels (χ^2^ = 3.43, *df* = 1, *P* = 0.06) with unsupplemented nestlings having lower hemoglobin levels than supplemented nestlings (Table 1). These results were likely because hemoglobin levels were negatively related to parasite abundance across parasite treatments (χ^2^ = 18.60, *df* = 1, *P* < 0.0001). The interaction between parasite and food treatment did not affect hemoglobin levels (χ^2^ = 1.29, *df* = 1, *P* = 0.26). Parasite treatment, food treatment, and the interaction between the two factors did not affect glucose levels (Tables 1 and S4) (parasite: χ^2^ = 0.41, *df* = 1, *P* = 0.52, food: χ^2^ = 0.83, *df* = 1, *P* = 0.36, interaction: χ^2^ = 0.00, *df* = 1, *P* = 0.96).

### 3.4 Immune responses of parasitized nestlings

Food treatment affected nestling antibody levels (Table S5) (GLMM, χ^2^ = 5.96, *df* = 1, *P* = 0.02; supplemented, *n* = 9: 0.53 ± 0.15, unsupplemented, *n* = 11: 0.24 ± 0.07) with supplemented nestlings having higher antibody levels than unsupplemented nestlings within the sham-fumigated treatment (Fig. 2A). Mean nestling antibody levels within a nest were negatively related to parasite abundance (Fig. 2B) (GLM, χ^2^ = 5.86, *df* = 1, *P* = 0.02). Food treatment did not affect haptoglobin levels (Table S5) (GLM, χ^2^ = 0.20, *df* = 1, *P* = 0.65; supplemented: 0.36 ± 0.06, unsupplemented: 0.32 ± 0.04; *n* = 5 nestlings for both treatments). Haptoglobin levels were not significantly correlated with antibody levels (GLM, χ^2^ = 0.39, *df* = 1, *P* = 0.53) or parasite abundance (GLM, χ^2^ = 0.08, *df* = 1, *P* = 0.78).

**Fig. 2.**
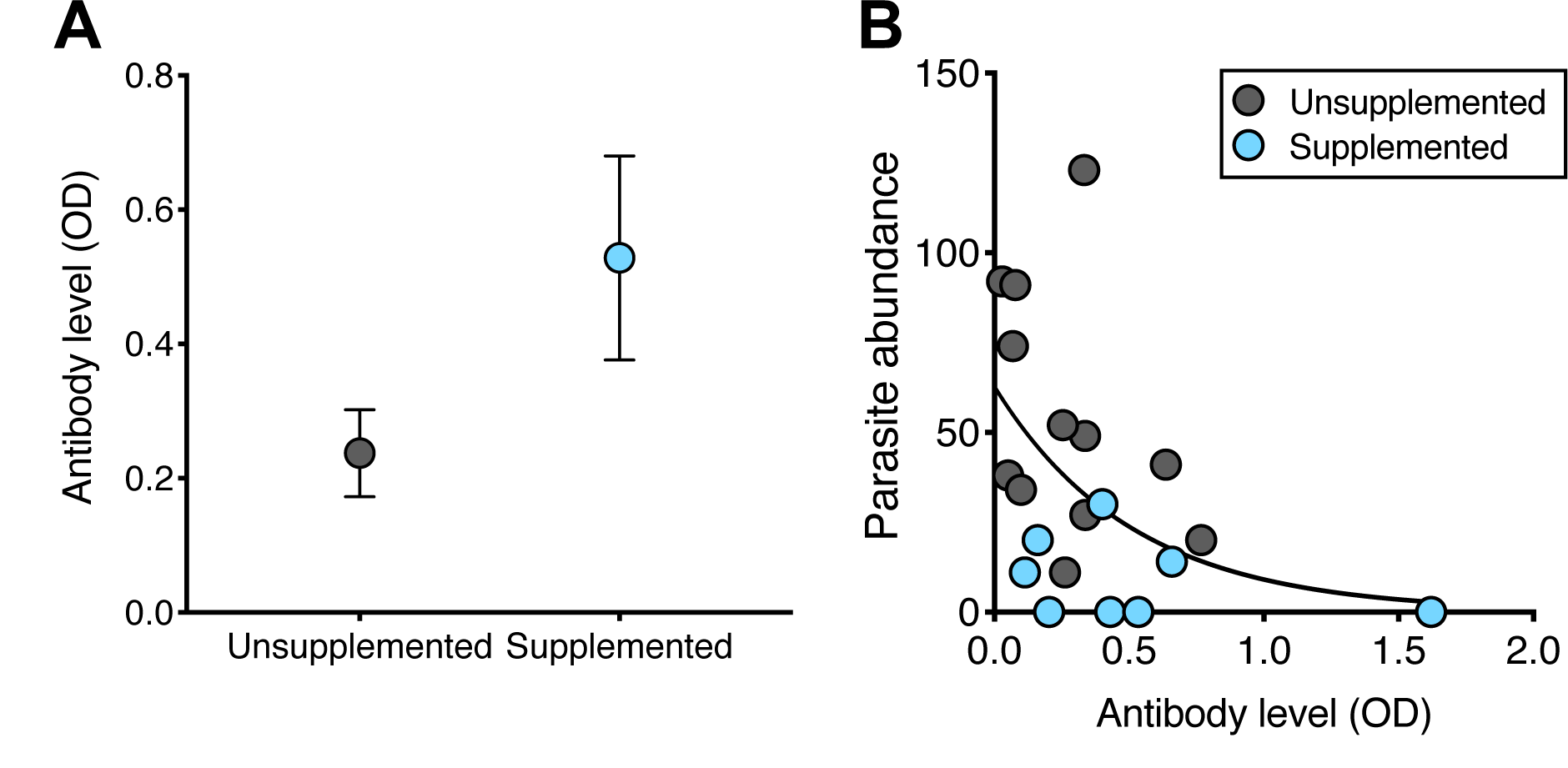
Within the parasitized treatment, supplemented birds had higher antibody levels than unsupplemented birds (A). Antibody levels were negatively related to parasite abundance (B).

### 3.5 Microbiota of parasitized nestlings

Bacterial community structure and membership differed between the parasitized nestlings and mealworms, but did not differ between food treatments for the nestlings (Fig. 3A and B) (PERMANOVA, structure: *F*_2,16_ = 2.15, *P* = 0.002, membership: *F*_2,16_ = 2.09, *P* = 0.001). Supplemented nestlings had higher bacterial diversity (sobs) compared to unsupplemented nestlings, but this difference was not significant for sobs (Fig. 4A) (Tables S5 and S6) (GLMM, χ^2^ = 3.07, *df* = 1, *P* = 0.08). Bacterial diversity (sobs) was not significantly related to antibody levels (Fig. 4B) (GLM, χ^2^ = 0.85, *df* = 1, *P* = 0.36) but negatively related to parasite abundance (Fig. 4C) (GLMM, χ^2^ = 4.17, *df* = 1, *P* = 0.04). Food treatment did not significantly affect the Shannon (GLMM, χ^2^ = 2.02, *df* = 1, *P* = 0.15) and Simpson index (Tables S5 and S6) (GLMM, χ^2^ = 1.41, *df* = 1, *P* = 0.24). Neither indices were significantly correlated with parasite abundance (GLMM, Shannon: χ^2^ = 0.29, *df* = 1, *P* = 0.59, Simpson: χ^2^ = 0.13, *df* = 1, *P* = 0.71) or antibody levels (GLM, Shannon: χ^2^ = 0.43, *df* = 1, *P* = 0.51, Simpson: χ^2^ = 0.30, *df* = 1, *P* = 0.59).

**Fig. 3.**
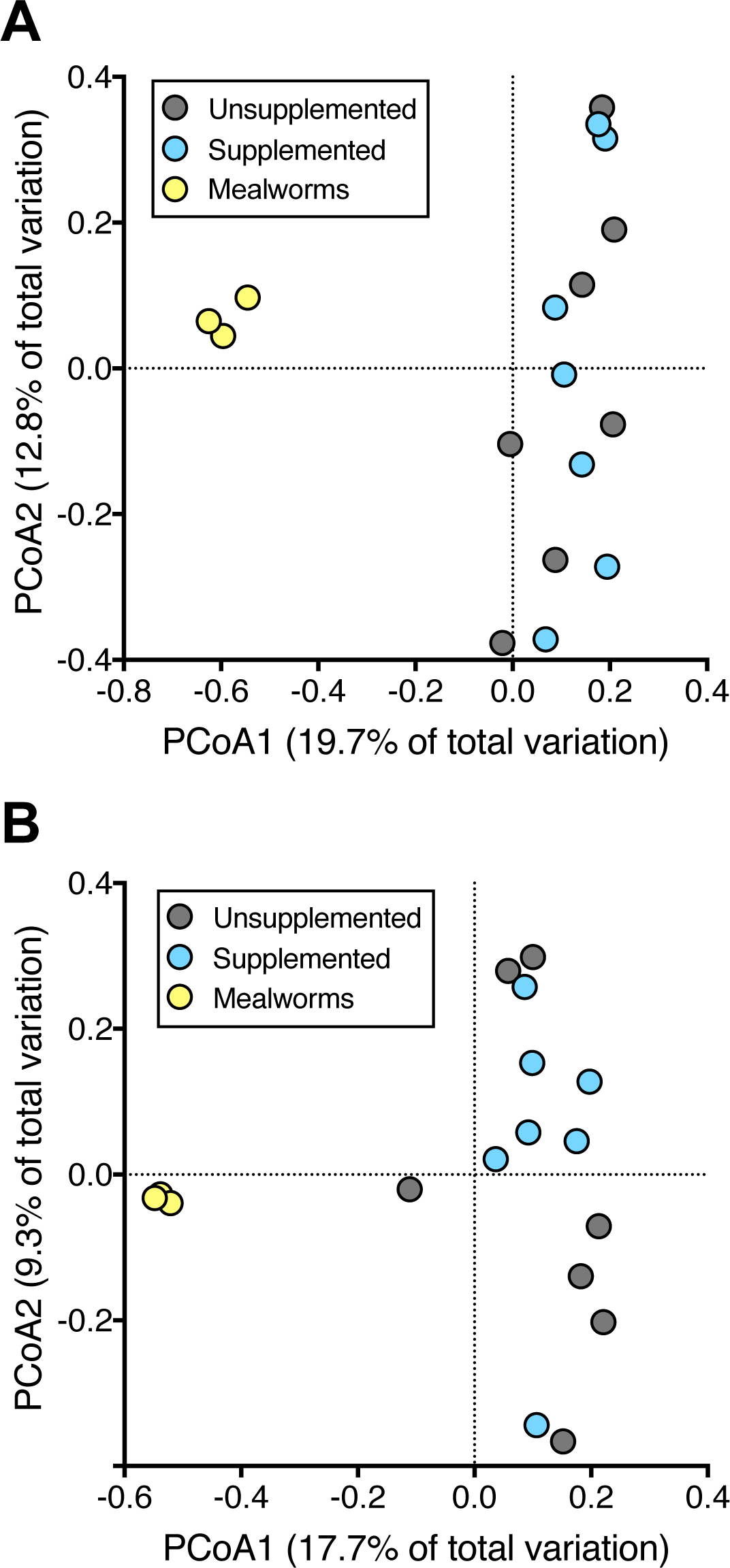
Within the parasitized treatment, bacterial community structure (A) and membership (B) of nestlings did not differ between food treatments. The bacterial community of mealworms was distinct from the bacterial community of the nestlings.

**Fig. 4.**
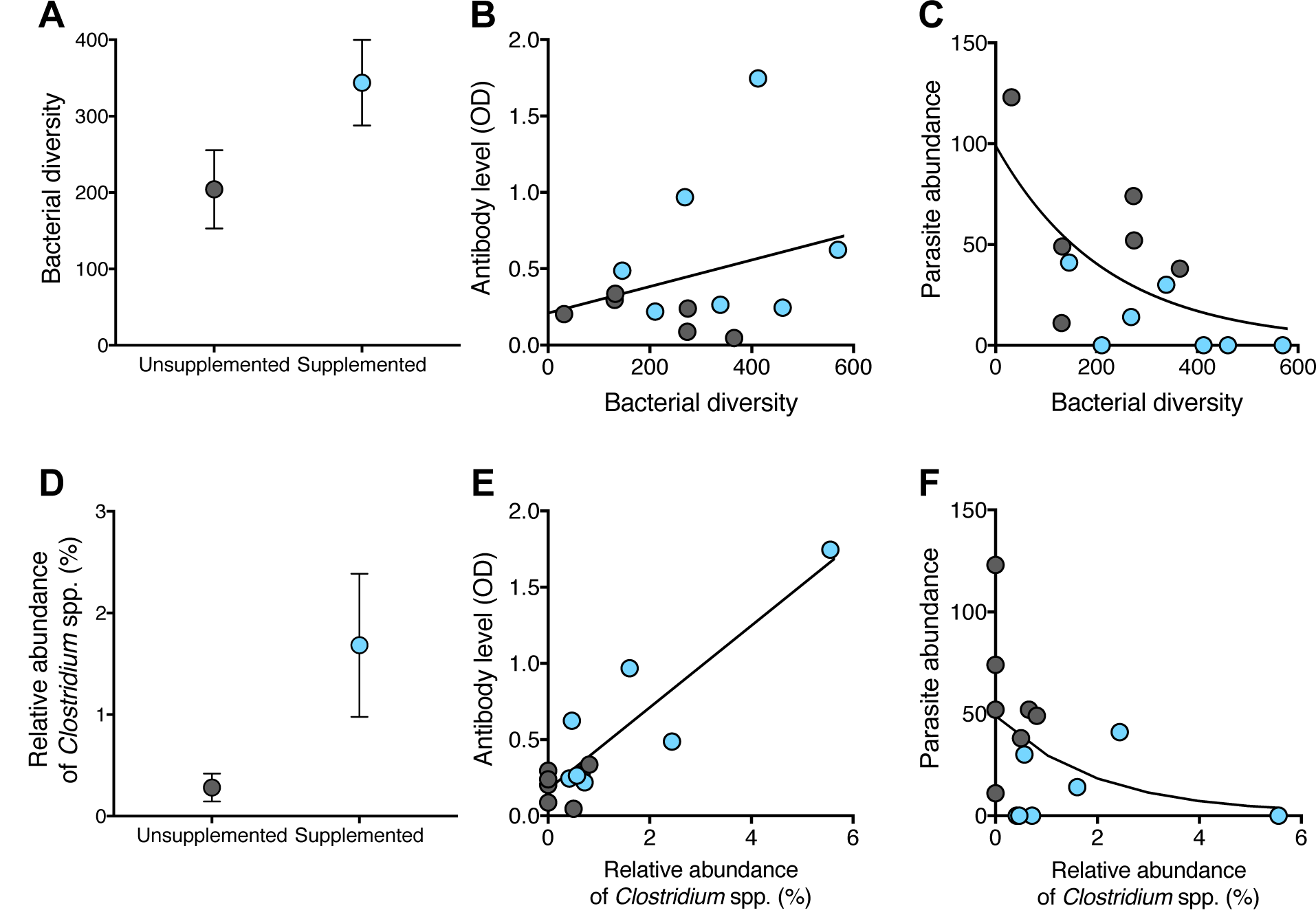
Effect of food treatment on microbiota of nestlings and the relationships among microbiota, immune response, and parasite abundance within the parasitized treatment. Supplemented nestlings had marginally higher bacterial diversity than unsupplemented nestlings (A). Bacterial diversity was not significantly related to antibody levels (B) but negatively related to parasite abundance (C). Relative abundance of *Clostridium* spp. was higher in supplemented nestlings than unsupplemented nestlings (D). *Clostridium* spp. abundance was positively related to antibody levels (E) and negatively related to parasite abundance (F).

The relative abundance of several bacterial phyla differed between the birds and mealworms. Birds in both treatments had higher relative abundances of phyla Planctomycetes (ANOVA, *F* = 8.61, *P* = 0.04) compared to the mealworms, but mealworms had higher relative abundances of phyla Cyanobacteria (ANOVA, *F* = 20.96, *P* = 0.001) and Tenericutes (ANOVA, *F* = 7.65, *P* = 0.04) than the birds (*P* < 0.05 for each pairwise test). Mealworms were dominated by phyla Firmicutes (36.03%), Proteobacteria (25.78%), Cyanobacteria (20.07%), Actinobacteria (10.14%), Tenericutes (5.25%), and Bacteroidetes (2.44%). Nestling feces were dominated by phyla Proteobacteria (28.24%), Actinobacteria (28.05%), Firmicutes (21.91%), Bacteroidetes (8.07%), Cyanobacteria (6.14%), Planctomycetes (1.47%), and Saccharibacteria (1.04%).

Several bacterial genera differed between the nestlings and mealworms (Table S7). Genera with known pathogenic bacterial species were then compared specifically between supplemented and unsupplemented birds. *Campylobacter spp., Listeria spp., Mycobacteria spp., Pasteurella spp., Salmonella spp., and Vibrios spp.* were not found in the feces of nestlings. Food treatment did not affect relative abundances of *Enterococcus spp., Escherichia spp., Lactobacillus spp., Staphylococcus spp., and Streptococcus spp*. Supplemented birds had higher relative abundances of genus *Clostridium* (Fig. 4D) (ANOVA, *F* = 7.92, *P* = 0.02) than unsupplemented birds. Relative abundance of *Clostridium spp.* correlated positively with antibody levels (Fig. 4E) (GLM, χ^2^ = 21.39, *df* = 1, *P* < 0.0001) and negatively with parasite abundance (Fig. 4F) (GLM, χ^2^ =3.59, *df* = 1, *P* = 0.058), but did not correlate significantly with haptoglobin levels (GLM, χ^2^ = 0.11, *df* = 1, *P* = 0.74).

## 4 DISCUSSION

The study showed that parasitism did not significantly affect growth or fledging success but negatively affected hemoglobin levels. Although nestlings maintained parasite tolerance across food treatments, food supplementation combatted the sublethal effects of parasitism by increasing nestling resistance to the parasites. However, food supplementation primarily benefited the birds (i.e. reduced parasite abundance) early in the breeding season. The extra food resources increased the nestling antibody response in parasitized nestlings, which decreased parasite abundance. Interestingly, haptoglobin levels did not relate to parasite abundance, nor were they affected by food supplementation, suggesting that these acute phase proteins are not related to blowfly resistance. Supplementation also increased gut bacterial diversity in parasitized nestlings, which was negatively related to parasite abundance, but not the antibody response. However, specifically, supplementation increased the relative abundance of genus *Clostridium* spp., which was positively related to the antibody response and negatively related to parasite abundance. The mealworms had a distinct bacterial community from the birds, which suggests that the nutritional composition of the mealworms influenced the gut microbiota of the birds. Overall, these results suggest that food supplementation facilitates an increase in gut *Clostridium* spp., which potentially primes the non-specific antibody response in nestlings to resist parasitic nest flies.

Without the extra food, bluebird nestlings are tolerant to the parasitic nest flies, at least in relation to survival (Grab *et al*. 2019). Therefore, why does food supplementation increase resistance when the birds are already relatively well-defended against the parasite? Parasitism negatively affected hemoglobin levels, which could have had lasting effects after the young left the nest. For example, blowflies did not affect nestling mass or fledging success of ovenbirds (*Seiurus aurocapilla*), however, fledgling survival and minimum distance traveled the first day after fledging was significantly lower when the ovenbirds were parasitized (Streby, Peterson & Kapfer 2009). Thus, when extra food is available resource-dependent resistance is likely beneficial to ameliorate the long-term or subtle adverse effects of parasitism.

Traditionally, food supplementation is thought to have a direct effect on the antibody response produced by the host. Immune function can be condition-dependent because the immune response can be energetically costly to produce and therefore only hosts in good condition may be physiologically able to invest in these responses (reviewed by Sheldon and Verhulst 1996; Lochmiller and Deerenberg 2000; Svensson *et al*. 1998). Other studies have found that supplemented nutrients can increase immunity (e.g. Ig antibodies) to parasites (Datta *et al*. 1998). For example, hosts fed a high-protein diet produced more IgG antibodies to parasitic worms compared to hosts fed a low-protein diet (Datta *et al*. 1998). Supplemented mealworms could provide the necessary additional protein to induce or increase the development of the IgY response in nestlings.

Food supplementation might have also or alternatively affected the nestling antibody response through changes in the gut microbiota. Host diet can affect the gut microbiota (David *et al*. 2014; Carmody *et al*. 2015; Knutie *et al*. 2017a). Specifically, my study found that food supplementation increased the relative abundance of *Clostridium* spp. and other studies have also found varying effects of host diet on *Clostridium* spp. abundance (e.g. in chickens) (Stutz & Lawton 1984; Mitsch *et al*. 2004; Jia *et al*. 2009). For example, Drew *et al*. (2004) found that protein supplementation increased *Clostridium perfringens* abundance. The high protein content of mealworms (46.44%) (Ravzanaadii *et al*. 2012) could be responsible for the increase *Clostridium spp.* abundance, but this idea requires further study.

A remaining question is whether the presence of *Clostridium spp.* in the gut is priming a non-specific antibody response to ectoparasitic nest flies. Pathogenic *Clostridium spp.* can activate the innate and adaptive immune system (including the IgY antibody response) in chickens (Kulkarni *et al*. 2007), which can be influenced by food supplementation (Yitbarek *et al*. 2012). Furthermore, priority effects, or the effect of one species on another species due to its earlier arrival, can influence the fitness of the multiple symbionts (Sousa 1992; de Roode *et al*. 2005; Devevey *et al*. 2015). For example, initial infection by parasitic worms can negatively affect the establishment of subsequent infections by other parasitic species in the host (Hoverman, Hoye & Johnson 2013; Wuerthner, Hua & Hoverman 2017), which could either be mediated by direct competition between the parasites or by the priming of the immune response of the host. Although gut bacteria and ectoparasites occupy different spaces on and in their host, the organisms are potentially exposed to the same circulating molecules related to the immune system (e.g. plasma IgY antibodies). However, future experimental studies are needed to determine whether increases in *Clostridium* spp. in hatchlings causally affects their IgY antibody response and resistance to *P. sialia*.

Between 1920-1970, bluebird populations declined throughout North America due, in part, to habitat destruction and the introduction of invasive species (Gowaty & Plissner 2015). However, in the 1980s, bluebird populations started to rebound because the public established nest boxes throughout the range of the bluebird. At study’s field site, I established 70 nest boxes in 2014-15, which increased to 150 boxes by 2017. Consequently, the number of nesting bluebird pairs increased from six pairs in 2015 to 31 pairs in 2017, and this number continues to increase. Across the past several decades, the public also became concerned about nest parasites affecting the health of the birds and therefore, have implemented methods to eliminate or deter the parasites (Zeleny 1976). Methods to remove parasites include removing old nests from the box (Møller 1989) and placing deterrents in the box, such as vanilla extract and insecticides (S.A.K. pers. comm.). My study suggests that supplying mealworms to bluebirds not only increases the health of nestlings, but also reduces parasite loads in the nests. However, these effects are most pronounced during the early part of the breeding season, likely because natural insect (food) availability is lowest at this time (Bowlin & Winkler 2004).

## 5 CONCLUSION

Determining the consequences of food supplementation is important because artificial bird feeding is a common activity for humans throughout the world (reviewed in Cox & Gaston 2018). The results of this study show that increasing food availability for hosts can decrease parasite pressure, especially during formative life stages. However, studies have shown that having an epicenter of feeders where animals are directly or indirectly interacting can increase transmission of more directly transmitted parasites (Becker *et al*. 2015). Therefore, the effect of food supplementation on parasite resistance could vary when considering how feeders affect parasite exposure. In this study, only the focal bluebirds were visiting the feeders (S.A.K. pers. obs.), which reduced contact with other birds. Therefore, when feeding breeding birds, especially box-nesting birds, placing individual feeders in the birds’ territory could increase host defenses without the risk of infection transmission.

Additionally, the results of the study might be particularly important for systems that are dealing with detrimental parasites, such as Darwin’s finches of the Galapagos Islands or pardalotes of Australia and their parasitic nest flies (Fessl *et al*. 2010; Edworthy, Langmore & Heinsohn 2019). When managing the parasites is not immediately possible, providing additional food resources to the host could help increase their resistance (or even tolerance; (Knutie *et al*. 2016; McNew *et al*. 2019)) to parasites. However, assessing the potential risks of providing additional resources to endangered or threatened species is required before attempting the method.

## Supporting information

Supplemental Results

## ACKNOWLEDGEMENTS

I thank Terry Whitworth for identifying *Protocalliphora sialia*, Steve Knutie and Doug Thompson for building nest boxes, and Itasca Biological Station (David Biesboer, Laura Domine, and Lesley Knoll) for logistical support. I also thank the following people for allowing us access to nest boxes on their property and their help in the field (in alphabetical order): Wade and Kelly Foy (Rock Creek General Store), Aaron Hebbeler, Jim Humeniuk, John and Dottie Hurlbert, Alvin and Dorothy Katzenmeyer, Lesley Knoll, Lake Itasca Pioneer Farmer’s, Helen and Ken Perry, Natasha, Jay, Tatiana, and Jesse Smrekar, and Doug and Dawn Thompson. I thank Brian Hiller, McKenzie Ingram, Allie Parker, and Darya Pokutnaya for field assistance, Lauren Albert for lab assistance, Sabrina McNew, Samantha Rumschlag, and Michael Mahon for help with the statistical analyses, and Kendra Maas at the University of Connecticut Microbial Analysis, Resources, and Services for workshop training and advice on the Mothur bioinformatics. The work was funded by a Savaloja Research Grant from the Minnesota Ornithologists’ Union, a Research Grant from the North American Bluebird Society, and start-up funds from the University of Connecticut. All applicable institutional guidelines for the care and use of animals were followed. The author declares no competing interests.

## DATA ACCESSIBILITY

Sequences have been uploaded to GenBank (BioProject accession number: available upon acceptance) and data have been uploaded to FigShare (doi available upon acceptance).

