## Supplemental Results for "Food supplementation affects gut microbiota and immunological resistance to parasites in a wild bird species"

S.A. Knutie

Table S1. Pairwise correlation coefficients between growth metrics (bill length, tarsus length, first primary feather length, and body mass).

| | Bill length | Tarsus length | Feather length | Body mass |
|---|---|---|---|---|
| Bill length | 1.00 | | | |
| Tarsus length | 0.37 | 1.00 | | |
| Feather length | 0.65 | 0.47 | 1.00 | |
| Body mass | 0.48 | 0.38 | 0.66 | 1.00 |

Table S2. The effect of parasite treatment, food treatment, and their interaction on parasite abundance.

| **Variables** | **Coefficients ± SE** |
|---|---|
| Parasite treatment | 12.89 ± 10.60 |
| Food treatment | 0.00 ± 10.60 |
| Parasite x food treatment | 42.66 ± 13.94 |

Table S3. The effect of food treatment, Julian date, and their interaction on parasite abundance within the sham-fumigated treatment.

| **Variables** | **Coefficients ± SE** |
|---|---|
| Food treatment | 21.24 ± 8.88 |
| Julian date | 0.05 ± 0.04 |
| Food treatment x Julian date | -0.11 ± 0.05 |

Table S4. The effect of parasite treatment, food treatment, and their interaction on growth, hemoglobin levels, and glucose levels.

| **Variables** | **Coefficients ± SE** |
|---|---|
| Bill length | |
| Parasite treatment | 0.00 ± 0.02 |
| Food treatment | -0.02 ± 0.02 |
| Parasite x food treatment | 0.01 ± 0.02 |
| Tarsus | |
| Parasite treatment | 0.02 ± 0.01 |
| Food treatment | 0.00 ± 0.01 |
| Parasite x food treatment | -0.01 ± 0.01 |
| First primary length | |
| Parasite treatment | 0.02 ± 0.06 |
| Food treatment | -0.10 ± 0.07 |
| Parasite x food treatment | 0.05 ± 0.08 |
| Body mass | |
| Parasite treatment | -0.00 ± 0.02 |
| Food treatment | -0.03 ± 0.02 |
| Parasite x food treatment | -0.00 ± 0.02 |
| Hemoglobin | |
| Parasite treatment | -0.05 ± 0.06 |
| Food treatment | -0.01 ± 0.07 |
| Parasite x food treatment | -0.10 ± 0.09 |
| Glucose | |
| Parasite treatment | -0.04 ± 0.08 |
| Food treatment | -0.05 ± 0.09 |
| Parasite x food treatment | 0.01 ± 0.12 |

Table S5. The effect of food treatment on immune responses and bacterial diversity metrics.

| **Variables** | **Coefficients ± SE** |
|---|---|
| IgY antibody levels | -0.28 ± 0.11 |
| Haptoglobin levels | -0.03 ± 0.07 |
| Bacterial diversity | |
| sobs | -0.30 ± 0.17 |
| Shannon index | -1.06 ± 0.74 |
| Simpson Index | -0.32 ± 0.27 |

Table S6. The effect of food treatment on mean ± SE bacterial diversity metrics. Numbers in the parentheses are the sample sizes of nests.

| **Variables** | Supplemented | Unsupplemented |
|---|---|---|
| sobs | 343.80 ± 56.04 (7) | 204.25 ± 51.28 (6) |
| Shannon index | 3.14 ± 0.41 (7) | 2.06 ± 0.64 (6) |
| Simpson index | 12.85 ± 3.38 (7) | 9.60 ± 5.00 (6) |

Table S7. Relative abundances (%) of bacterial genera that differ between the birds (supplemented: S, unsupplemented: US) and mealworms (MW). Numbers (%) in bold indicate which treatment of birds are statistically different from the mealworms (Tukey: $P < 0.05$).

| Genus | MW | S | US | Statistics |
|---|---|---|---|---|
| *Brachybacterium* | 1.90 | **0.08** | **0.00** | $F = 67.12$, $P < 0.0001$ |
| *Brevibacterium* | 5.08 | **0.13** | **0.00** | $F = 103.35$, $P < 0.0001$ |
| *Candidatus* | 0.00 | **2.68** | **2.41** | $F = 18.38$, $P = 0.002$ |
| *Dietzia* | 1.25 | **0.00** | **0.00** | $F = 11.53$, $P = 0.01$ |
| *Erysipelatoclostridium* | 0.00 | **2.46** | **3.89** | $F = 9.17$, $P = 0.03$ |
| *Hyphomicrobium* | 0.00 | **1.43** | **0.97** | $F = 8.14$, $P = 0.04$ |
| *Kocuria* | 1.90 | **0.20** | **0.00** | $F = 34.96$, $P = 0.0001$ |
| *Lactobacillus* | 8.96 | **0.45** | **0.00** | $F = 31.82$, $P = 0.0001$ |
| *Lactococcus* | 15.88 | **0.72** | **0.67** | $F = 31.91$, $P = 0.0001$ |
| *Methylobacterium* | 0.62 | **3.83** | **3.94** | $F = 9.27$, $P = 0.03$ |
| *Mycobacterium* | 0.00 | **2.53** | **2.50** | $F = 11.55$, $P = 0.01$ |
| *Nocardioides* | 0.00 | **6.01** | **5.93** | $F = 24.47$, $P < 0.001$ |
| *Pantoea* | 1.90 | **0.66** | **0.66** | $F = 7.57$, $P = 0.04$ |
| *Parabacteroides* | 1.27 | **0.08** | **0.00** | $F = 7.56$, $P = 0.04$ |
| *Parasutterella* | 1.90 | **0.00** | **0.00** | $F = 40015.68$, $P < 0.0001$ |
| *Rhizobium* | 0.00 | **1.40** | **1.67** | $F = 9.85$, $P = 0.02$ |
| *Rhodococcus* | 0.00 | **2.32** | **2.13** | $F = 9.77$, $P = 0.02$ |
| *Spiroplasma* | 10.78 | **0.33** | **0.20** | $F = 64.65$, $P < 0.0001$ |
| *Staphylococcus* | 5.07 | **0.52** | **0.35** | $F = 11.96$, $P = 0.01$ |
| *Variibacter* | 0.00 | **0.97** | 0.56 | $F = 7.75$, $P = 0.04$ |
| *Weissella* | 9.41 | **0.22** | **0.20** | $F = 35.30$, $P = 0.0001$ |
| *Xanthomonas* | 1.90 | **0.08** | **0.18** | $F = 18.99$, $P = 0.001$ |

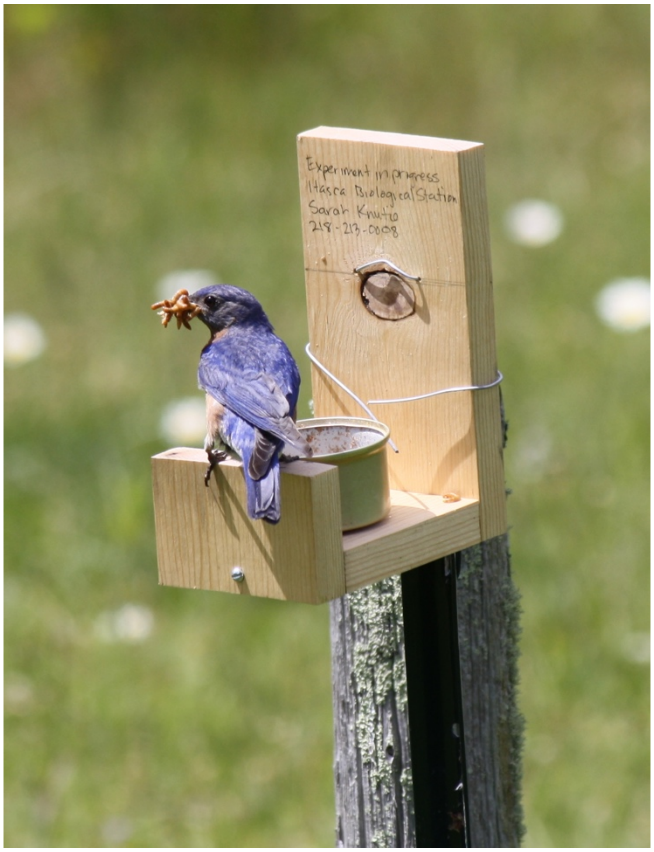


Fig. S1. Experimental mealworm feeder being used by a male eastern bluebird.
